# Mechanism of Interaction of BMP and Insulin Signaling in *C. elegans* Development and Homeostasis

**DOI:** 10.1101/777805

**Authors:** James F. Clark, Emma J. Ciccarelli, Peter Kayastha, Gehan Ranepura, Muhammad S. Hasan, Alicia Meléndez, Cathy Savage-Dunn

## Abstract

A small number of peptide growth factor ligands are used repeatedly in development and homeostasis to drive programs of cell differentiation and function. Cells and tissues must integrate inputs from these diverse signals correctly, while failure to do so leads to pathology, reduced fitness, or death. Previous work using the nematode *C. elegans* identified an interaction between the bone morphogenetic protein (BMP) and insulin/IGF-1-like signaling (IIS) pathways in the regulation of lipid homeostasis. The molecular components required for this interaction, however, were not known. Here we report that INS-4, one of 40 insulin-like peptides (ILPs), is regulated by BMP signaling to modulate fat accumulation. Furthermore, we find that the IIS transcription factor DAF-16/FoxO, but not SKN-1/Nrf, acts downstream of BMP signaling in lipid homeostasis. Interestingly, BMP activity alters sensitivity of these two transcription factors to IIS-promoted cytoplasmic retention in opposite ways. Finally, we probe the extent of BMP and IIS interactions by testing two additional IIS functions, dauer formation and autophagy induction. Coupled with our previous work and that of other groups, we conclude that BMP and IIS pathways have at least three modes of interaction: independent, epistatic, and antagonistic. The molecular interactions we identify provide new insight into mechanisms of signaling crosstalk and potential therapeutic targets for IIS-related pathologies such as diabetes and metabolic syndrome.

## Introduction

Cell signaling pathways are integral to normal development and homeostasis, and their misregulation is frequently associated with pathological conditions including cancer, cardiovascular disease, and metabolic syndrome. Cells and tissues have distinctive patterns of responsiveness to a variety of secreted peptide ligands. The correct integration of these signals allows for robust control of development and sensitive regulation of homeostatic processes. We have previously identified a regulatory interaction between bone morphogenetic protein (BMP) and insulin/IGF-1-like signaling (IIS) pathways in the regulation of lipid stores in the nematode *Caenorhabditis elegans*. BMPs are members of the Transforming Growth Factor beta (TGFβ) family of peptide ligands. BMPs are conserved across animal phyla, and their actions are traditionally known for roles in development, growth, and differentiation (Shi and Massague 2003; Wu and Hill 2009). Our research, however, has explored the homeostatic effects of BMP signaling, using the powerful genetic tractability and imaging tools available in *C. elegans* to elucidate interactions between signaling pathways. In addition to our work, roles for TGFβ family members in regulating metabolic homeostasis in vertebrates have been elucidated recently (Wang et al. 2004; Fain et al. 2005; Sjoholm et al. 2006; Bottcher et al. 2009; Shen et al. 2009). BMPs in particular may play a role in insulin regulation and age-related insulin resistance (Chattopadhyay et al. 2017). Unlike BMP signaling, IIS is well-known to function in homeostatic processes. In mammals, IIS regulates glucose uptake and metabolic homeostasis (Anisimov and Bartke 2013). Disruptions to insulin balance, such as insulin resistance, can lead to multiple metabolic disorders and diseases, including obesity and type II diabetes. Studying these pathways in *C. elegans* is fruitful due to the high degree of conservation of the respective signaling pathways.

The BMP signaling pathway in *C. elegans* includes founding members of the Smad family of signal transducers, SMA-2, SMA-3, and SMA-4 (Savage et al. 1996; Savage-Dunn and Padgett 2017). DBL-1, the *C. elegans* BMP2/4 homolog, plays a major role in body size regulation, male tail development, and mesodermal patterning (Suzuki et al. 1999; Foehr et al. 2006). Initial identification of several genes related to fat metabolism through microarray analysis in a *dbl-1* mutant background (Liang et al. 2007) led to the hypothesis that DBL-1/BMP is a regulator of metabolism. We confirmed this hypothesis, showing that disruptions to DBL-1 signaling lead to reduced lipid stores and altered lipid droplet morphology. Additionally, we determined that this regulation of lipid storage by DBL-1/BMP is dependent on IIS activity (Clark et al. 2018).

In *C. elegans*, the IIS pathway uses a single insulin receptor DAF-2/InsR (Gottlieb and Ruvkun 1994; Kimura et al. 1997) in conjunction with 40 insulin-like peptides (ILPs) to regulate multiple homeostatic functions through the control of transcription factors, such as DAF-16/FoxO (Ogg et al. 1997; Paradis and Ruvkun 1998; Lin et al. 2001) and SKN-1/Nrf (Tullet et al. 2008).

DAF-2/IIS has prolific effects on the development and homeostasis of the worm; disruptions to DAF-2/IIS lead to phenotypes in dauer formation, longevity, stress tolerance, innate immunity, germline maintenance, metabolism, and autophagy (Gems et al. 1998; Tissenbaum and Ruvkun 1998; Melendez et al. 2003; Jia et al. 2004; Brys et al. 2010; Michaelson et al. 2010; Kaplan and Baugh 2016).

In this study, we show an antagonistic effect between DBL-1/BMP and DAF-2/IIS in a *daf-2* mutant background when considering dauer formation and autophagy induction. Together with our previously published data on lipid stores and body size (Clark et al. 2018), our results outline three potential modes of interaction between DBL-1/BMP and DAF-2/IIS: independent, as in body size; epistatic; as in lipid regulation; and antagonistic, as seen in dauer formation and autophagy induction. We observe that Smad signaling in the hypodermis (epidermis) regulates expression of *ins-4*, and that *ins-4* mutants express a high-fat phenotype similar to that of *daf-2*, implicating INS-4 as a downstream mediator. Additionally, we find that loss of *dbl-1* alters the subcellular localization of DAF-16::GFP and SKN-1::GFP, and adjusts their sensitivity to DAF-2/IIS in opposite directions. Taken together, our data identify new methods of interaction that mesh the signaling outputs of two well-conserved pathways, BMP and IIS, in the regulation of homeostatic functions.

## Materials and Methods

### Nematode strains and growth conditions

*C. elegans* were maintained on *E. coli* (DA837) at 15°C, 20°C, and 25°C as specified. The wild-type strain used in this study was N2. The strains used in this study are as follows: LT121 *dbl-1(wk70)*, CB678 *lon-2(e678)*, CS24 *sma-3(wk30)*, CB1370 *daf-2(e1370)*, RB2544 *ins-4(ok3534)*, HT1693 *unc-119(ed3);wwEx63*, DA2123 *adIs2122*, MAH14 *daf-2(e1370);adIs2122*, TJ356 *zIs356[daf-16::gfp]*, LD1 *ldIs7[skn-1::gfp]*, CS683 *daf-2(e1370)sma-3(wk30)*, CS681 *daf-2(e1370);lon-2(e678)*. Crosses were used to obtain: CS663 *sma-3(wk30);ins-4(ok3534)*, CS633 *sma-3(wk30);wwEx63*, CS661 *daf-2(e1370);dbl-1(wk70);adIs2122*, CS634 *daf-2(e1370);lon-2(e678);adIs2122*, CS626 *dbl-1(wk70);adIs2122*, CS627 *lon-2(e678);adIs2122*, CS685 *dbl-1(wk70);zIs356*, and CS679 *dbl-1(wk70);ldIs7*.

### INS-4p::GFP Expression

Animals were grown at 20°C and imaged as day 1 adults. Images were taken on a Leica Microsystems Confocal microscope using a 20X objective. Camera settings were maintained identically between all samples and trials. Compound images were produced from compiling Z-stack images. n>10 per strain repeated in triplicate.

### Oil Red O staining

Protocol adapted from (O’Rourke et al. 2009). Animals were collected at the L4 stage in PCR tube caps and washed three times in PBS. Worms were then fixed for 1 hour in 60% isopropanol while rocking at room temperature. The isopropanol was removed and worms were stained overnight with 60% Oil Red O solution while rocking at room temperature. Oil Red O was removed and the worms were washed once with PBS w/ .01% Triton and left in PBS. Worms were mounted and imaged using an AxioCam MRc camera with AxioVision software. Images were taken using a 40X objective. Oil Red O stock solution was made with 0.25g Oil Red O in 50mL isopropanol. Intensity of the post-pharyngeal intestine was determined using ImageJ software. Pixel intensity was measured in the green color channel of the images. Three measurements using a fixed size area were taken for each worm with background intensity subtracted for each individual picture. Statistical comparisons (two-way ANOVA, one-way ANOVA with post-hoc Tukey’s multiple comparisons test, and unpaired t-test) were performed using GraphPad Prism 7 software. For each experiment, n>20 per strain repeated in triplicate.

### Dauer formation assay

Five to ten L4 animals were placed per plate to lay eggs overnight at 15°C. Adults and L1s were removed on the subsequent day with M9. Plates with remaining eggs were then placed at either 20°C or 25°C to develop. Animals were then observed 48 hours later; dauer-like larvae were tallied versus L4s and adults. Statistical comparisons (unpaired t test and two-way ANOVA) were performed using GraphPad Prism 7 software. n>50 per strain at each temperature, repeated in triplicate.

### GFP::LGG-1 autophagy assay

Autophagy was assayed via the formation of GFP punctae as described in (Palmisano and Meléndez 2016). Animals were grown at 20°C and imaged at the L3 larval stage; dauer larvae were not imaged or quantified. GFP::LGG-1 punctae were counted in the seam cells of the hypodermis. Images were taken using a Zeiss ApoTome with AxioVision software and a 100X objective. Exposure times were kept consistent across strains within each experiment. Statistical comparisons (unpaired t test and one-way ANOVA) were performed using GraphPad Prism 7 software. n=15 per strain, repeated in duplicate.

### DAF-16::GFP and SKN-1::GFP Localization Assays

RNAi induction and feeding were carried out as described in (Kamath et al. 2000; Liang et al. 2013). For DAF-16::GFP, L4 animals were placed on EZ worm plates supplemented with 1mM IPTG and 50mg/ml carbenicillin overnight. Adults were moved to new RNAi plates and allowed to lay eggs for 4-6 hours to synchronize the progeny. Animals were then allowed to develop at 15°C until the L3 larval stage followed by a shift to 20°C to bypass the dauer induction of *daf-2* knockdown. L4 animals were assayed for GFP localization. L4440 was used as the empty vector control. Images were taken using a Zeiss ApoTome with AxioVision software and a 40X objective. Exposure times were kept consistent across strains within each experiment. GFP localization was categorized as none, low, medium, or high. n>20 per strain, repeated in duplicate. For SKN-1::GFP, L4 animals were placed on EZ worm plates supplemented with 1mM IPTG and 50mg/ml carbenicillin seeded with HT115 *E. coli*. Animals were allowed to lay eggs overnight. Adults and L1 larvae were removed the following day with M9. Remaining eggs were synchronized with a 4 to 6 hour hatch, with the L1s being transferred to new RNAi plates. Animals were then allowed to develop at 15°C until the L3 larval stage followed by a shift to 25°C to bypass the dauer induction of *daf-2* knockdown. Animals were left at 25°C overnight and then assayed for GFP localization. L4440 was used as the empty vector control. Images were taken using a Zeiss ApoTome with AxioVision software and a 20X objective. Exposure times were kept consistent across strains within each experiment. GFP localization was categorized as low, medium, or high. n>40 per strain, repeated in duplicate. Statistical comparisons (chi-square test) were performed using GraphPad Prism 7 software.

## Results

### *ins-4* Regulation in the Hypodermis by SMA-3/Smad

In previous work, we showed that SMA-3/Smad activity in the hypodermis (epidermis) regulates fat accumulation in the *C. elegans* intestine, and that SMA-3/Smad regulation of fat stores depends on the DAF-2 Insulin Receptor (InsR) (Clark et al. 2018). We therefore hypothesized that SMA-3/Smad may regulate expression of an insulin-like peptide (ILP) in the hypodermis, which then modulates DAF-2/InsR activity. To identify such an ILP, we considered two transcriptional targets previously identified by microarray analysis, *ins-4* and *ins-7* (Liang et al. 2007). Of these, *ins-4* is reported to be expressed in hypodermis and neurons, whereas *ins-7* expression has been observed in the intestine and in neurons (Murphy et al. 2007; Ritter et al. 2013). To observe INS-4 regulation, we obtained an *ins-4p::gfp* reporter and crossed it into a *sma-3* mutant background. In wild-type animals carrying the *ins-4p::gfp* reporter, GFP was visible in neurons while only faint GFP expression was observed in hypodermis. However, in *sma-3;ins-4p::gfp* animals, GFP expression was starkly increased in the hypodermis, indicating an increase in *ins-4* expression (Figure 1A). This observation indicates that DBL-1/BMP signaling normally down-regulates the expression of *ins-4*, corroborating the data from the microarray analysis. We conclude that SMA-3/Smad regulates expression of *ins-4* in the hypodermis.

**Figure 1.**
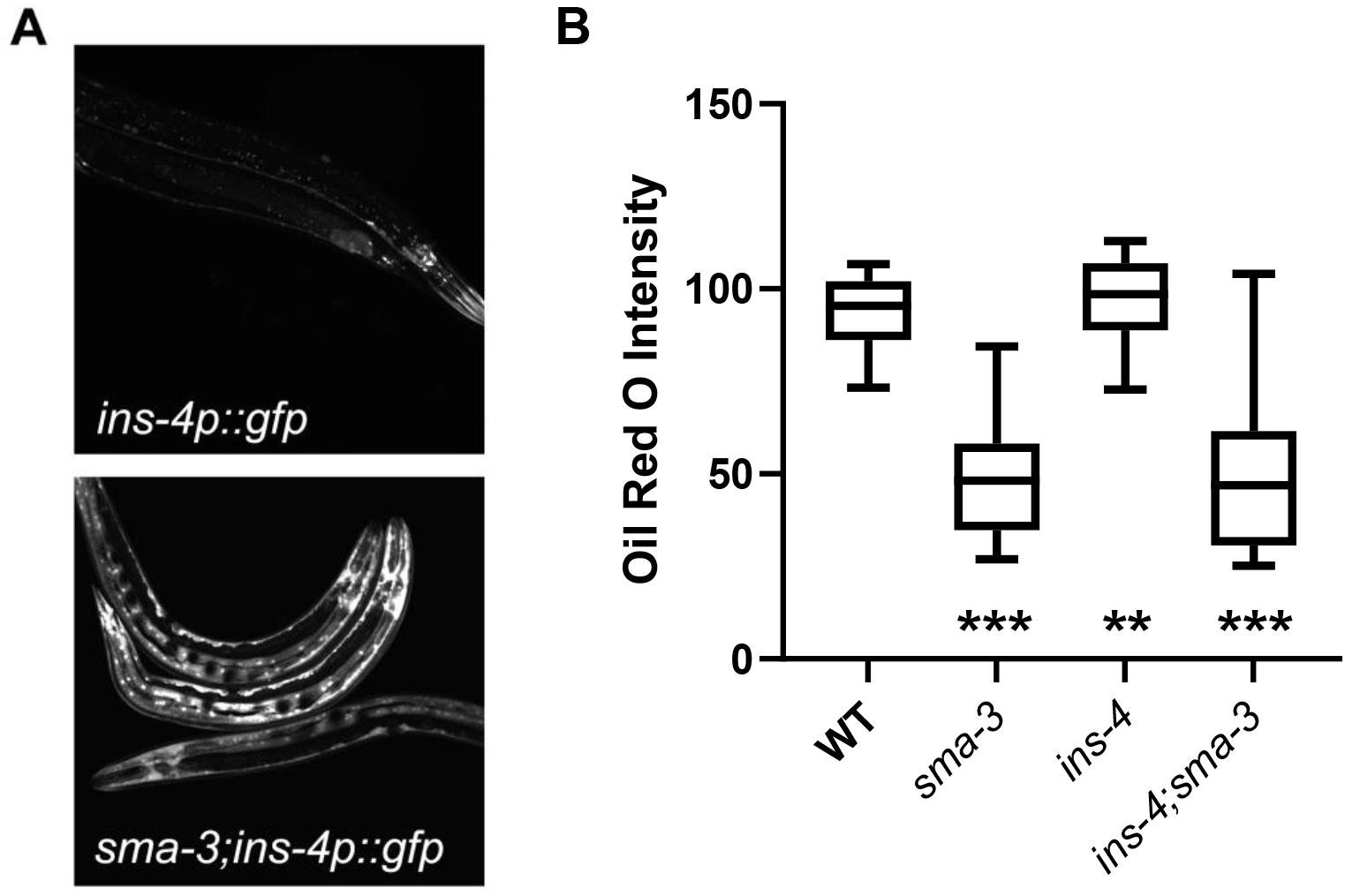
INS-4 acts downstream of SMA-3 and regulates lipid metabolism. A) Images of animals expressing *ins-4p::*GFP. Control animals displayed *ins-4p::*GFP in head neurons, with faint expression in the hypodermis. Upon loss of *sma-3, ins-4p::*GFP expression is significantly increased in the hypodermis. Images taken at 200X, camera settings were kept constant between all strains. B) Loss of *ins-4* results in an increase in intestinal lipid stores via Oil Red O staining. Animals were grown at 20°C and stained at the L4 larval stage. Quantification was done for equivalent regions of the intestine just posterior to the pharynx. For all graphs, asterisks across the bottom denote significance compared to Control, n.s. not significant, * p value < 0.05, *** p value <0 .001, boxes denote the 2^nd^ and 3^rd^ quartiles, whiskers denote min and max values.

### Loss of *ins-4* Results in a High-fat Phenotype

If INS-4 acts as a mediator between Smad signaling in the hypodermis and DAF-2/InsR signaling in the intestine, it may be required for normal lipid accumulation. To date, single mutants for *C. elegans* ILP genes have not displayed mutant phenotypes, due to redundancy between the genes (Ritter et al. 2013; Zheng et al. 2018a). Nevertheless, we measured lipid accumulation in *ins-4* mutant animals via Oil Red O staining. We observed an increase of ∼20% in the *ins-4* mutants compared to wild-type (p=0.0016) (Figure 1B,C), similar to the increase seen in *daf-2* mutants (Kimura et al. 1997; Clark et al. 2018). We therefore conclude that INS-4 represses fat accumulation and is required for normal lipid stores. These results indicate that the loss of *ins-4* is sufficient to alter significantly the level of lipids in the intestine. As previously published, we saw a significant decrease in *sma-3* animals by ∼42% (p<0.0001). To test the interaction between *sma-3* and *ins-4*, we generated a *sma-3;ins-4* double mutant and determined its phenotype. *sma-3;ins-4* double mutants have a low fat accumulation phenotype compared to wild type (p<0.0001) (Figure 1B,C), suggesting the involvement of additional ILPs in this interaction.

### DBL-1/BMP Signaling Promotes Dauer Arrest in a DAF-2/IIS-Sensitized Background

Since we discovered that DBL-1/BMP and DAF-2/IIS pathways interact to regulate lipid accumulation, we sought to characterize the relationship between the DBL-1/BMP and DAF-2/IIS pathways in other biological contexts. During development, animals exposed to harsh environmental conditions prior to the third larval stage can activate the stress-adaptive and growth-arrested dauer larval program. IIS is one of the primary regulators of the dauer decision, and *daf-2(e1370)* mutants have a temperature-sensitive dauer-constitutive phenotype. We investigated the degree of dauer formation in animals containing mutations in BMP and IIS pathways at the restrictive temperature, 25°C, and the semi-permissive temperature, 20°C. Normally, the DBL-1/BMP pathway has no effect on dauer, as *dbl-1* single mutants, and other mutants in the pathway, have no observable dauer phenotype at any temperature. As expected, when we placed *sma-3* or *lon-2* mutants at 25°C, we observed no dauer arrest phenotype, as in our wild-type control. When *daf-2* single mutants were placed at the restrictive temperature, we observed dauer arrest in 98.4±0.7% of the progeny, similar to previously published data. This phenotype was also observed in the *daf-2;sma-3* and *daf-2;lon-2* double mutants, with 99.5±0.6% and 97.6±1.7% dauer formation, respectively.

When animals were grown at the semi-permissive temperature, 20°C, however, DBL-1/BMP signaling had an observable impact on arrested animals in the *daf-2* mutant background. Again, the *sma-3* and *lon-2* mutants and wild type had no dauer induction at 20°C. *daf-2* single mutants exhibited the dauer arrest phenotype in 54.5±6.3% of the progeny at the semi-permissive temperature. The *daf-2;sma-3* double mutants, in contrast, exhibited the dauer arrest phenotype in only 23.9±4.6% of the animals, while the *daf-2;lon-2* double mutants exhibited the dauer arrest phenotype in 73.4±7.3% of the animals (Table 1). Two-way ANOVA indicates a significant (p=0.0006) interaction between the two pathways. This result indicates that DBL-1/BMP signaling has a significant effect on the dauer arrest phenotype in the DAF-2/IIS-sensitive background. DBL-1/BMP signaling appears to modulate dauer formation in a dose-dependent manner, where reduced signaling, as in *daf-2;dbl-1* double mutants, reduces the occurrence of dauer-like animals, while increased signaling, as in *daf-2;lon-2* mutants, increases the occurrence of dauer-like animals.

**Table 1.** DBL-1 Pathway Modulates Dauer Constitutive Phenotype of *daf-2* Mutants at 20°C. In a WT background, DBL-1 signaling exerts no visible effect on dauer induction at both 20°C and 25°C. However, in a DAF-2-sensitized background, DBL-1 signaling promotes dauer arrest at 20°C. This effect is lost when observed at 25°C, where the loss of DAF-2 signaling is most potent. Values given are percent arrested dauer-like larva of total progeny ± SEM.

| Strain | 20°C |  | 25°C |  |
| --- | --- | --- | --- | --- |
|  | % Arrested | N | % Arrested | N |
| N2 | 0 | 239 | 0 | 228 |
| *sma-3(wk30)* | 0 | 243 | 0 | 178 |
| *lon-2(e678)* | 0 | 269 | 0 | 214 |
| *daf-2(e1370)* | 54.5 ± 4.5 | 273 | 98.4 ± 0.5 | 236 |
| *daf-2(e1370);sma-3(wk30)* | 23.9 ± 3.3 | 200 | 99.6 ± 0.4 | 221 |
| *daf-2(e1370);lon-2(e678)* | 73.5 ± 5.2 | 317 | 97.6 ± 1.2 | 199 |

### DBL-1/BMP Signaling Enhances Autophagic Induction in a DAF-2/IIS-Sensitized Background

Autophagy is a cellular process involved in the recycling of cellular debris and other material, and is vital for the maintained health of an organism (Melendez and Levine 2009). As a known regulator of autophagy, DAF-2/IIS acts to suppress the induction of autophagosome formation, with *daf-2* mutants exhibiting increased levels of autophagy. Additionally, autophagy is required for proper dauer development in *daf-2* mutants (Meléndez et al. 2003). DBL-1/BMP signaling has also been shown to inhibit autophagy, as mutants in the BMP pathway elevate clearance of aggregates of SQST-1/p62 autophagy receptor induced by proteotoxic stress (Guo et al. 2014). We reproduced the effect of BMP signaling on SQST-1::GFP accumulation following RNAi inhibition of SMA-3/Smad (S1 Fig).

We also measured autophagosome levels through the use of GFP::LGG-1, an autophagosome reporter (Palmisano and Meléndez 2016). LGG-1/LC3 is a protein involved in the formation, elongation, and fusion of autophagosomes (Meléndez et al. 2003). Normally localized in a diffuse cytosolic pattern, when autophagic conditions are induced, GFP::LGG-1 becomes localized to pre-autophagosomal structures and autophagosomes. The localization of GFP::LGG-1 has been previously observed in hypodermal seam cells. The seam cells form a row of cells that run the length of the hypodermis on each lateral side of the animal, and undergo remodelling in the development of dauer animals. Thus, seam cells were observed due to their size and ease in imaging. *daf-2* mutants are known to show increased levels of autophagy when grown at the semi-permissive and restrictive temperatures, 20°C and 25°C respectively. To determine if DBL-1/BMP signaling would behave similarly in the regulation of autophagy as it did with dauer formation, we grew the animals at 20°C. We observed an average of 5.51±2.42 punctae per seam cell in *daf-2* single mutants at 20°C. In *daf-2;dbl-1* double mutants, the average was decreased to 3.85±2.20 (p<0.0001), while in *daf-2;lon-2* double mutants, the average was increased to 6.90±2.75 (p<0.0001) (Figure 2). Therefore, DBL-1/BMP activity antagonizes the effect of DAF-2/IIS in the induction of autophagy.

**Figure 2.**
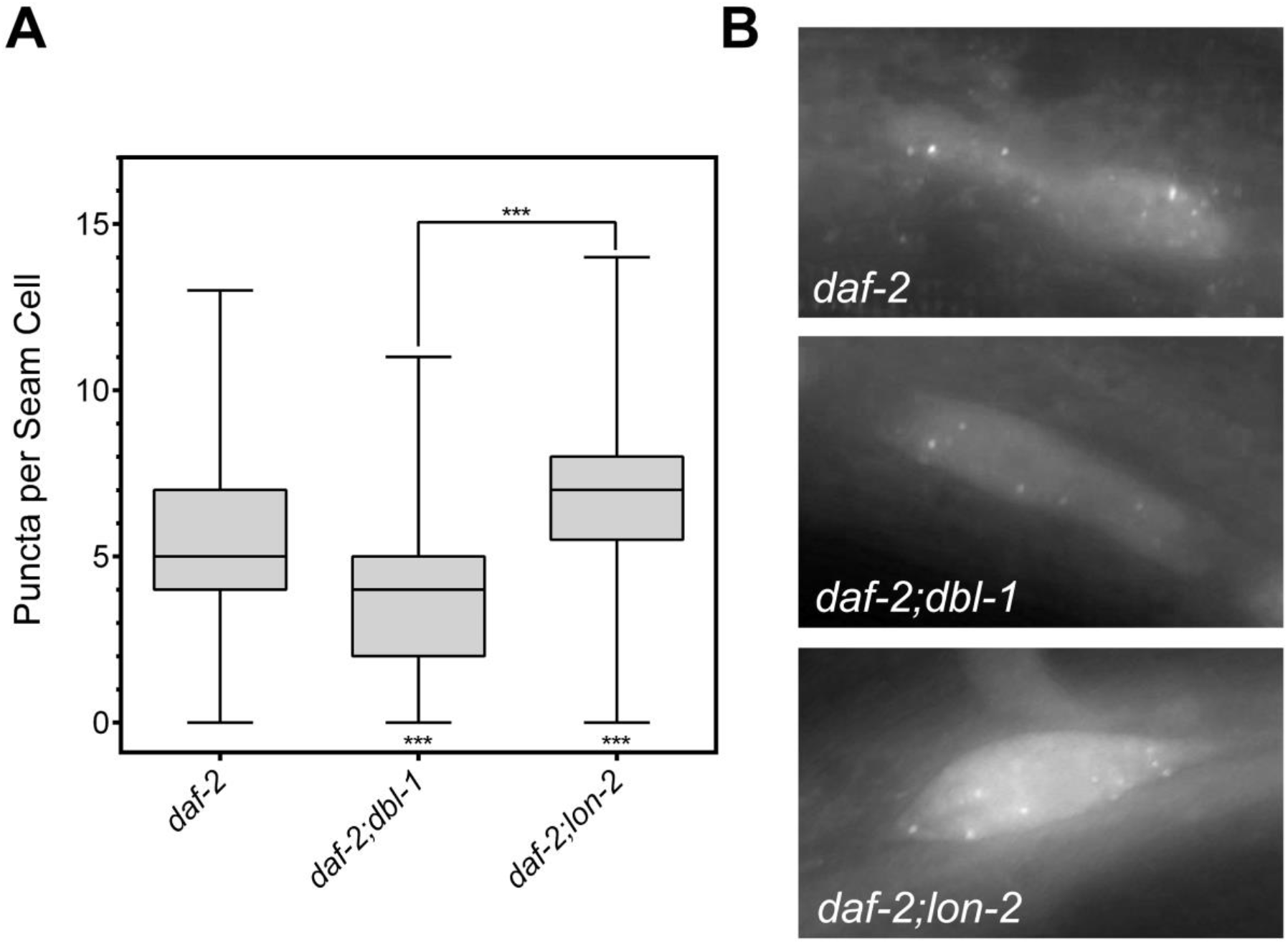
DBL-1 signaling promotes autophagy in a DAF-2-sensitized background. A) When autophagy is induced, via a reduction in DAF-2 signaling, DBL-1 signaling promotes autophagy, as *daf-2(e1370);dbl-1(wk70)* mutants show a reduction in GFP puncta, while *daf-2(e1370);lon-2(e678)* mutants show an increase in puncta. B) Images of GFP::LGG-1 localization in seam cells of DBL-1 pathway mutants under autophagic induction via *daf-2* mutant background at 630X. All animals were grown at 20°C and imaged at the L3 larval stage. *** p value < 0.001, boxes denote 2^nd^ and 3^rd^ quartiles, whiskers denote min and max values.

The observations presented above, combined with previous work from our lab (Clark et al. 2018), identifies three modes of interaction between DBL-1/BMP signaling and DAF-2/IIS: independent, epistatic, and antagonistic. DBL-1/BMP and DAF-2/IIS regulate body size independently, as double mutants in the two pathways were observed to have additive effects (Clark et al. 2018). Epistatic effects were observed in the regulation of lipids with the high-fat phenotype of *daf-2* mutants completely masking the low-fat phenotype of *sma-3* in *daf-2;sma-3* double mutants. Lastly, the data observed in this study reveals a third relationship between the two pathways; DBL-1/BMP signaling appears to antagonistically affect DAF-2/IIS phenotypes.

However, these effects are only observed in a DAF-2/IIS-compromised background. To elucidate further the context-dependent mechanisms by which DBL-1/BMP interacts with DAF-2/IIS, we explored interactions with transcription factors downstream of DAF-2.

### Loss of DBL-1/BMP Signaling Alters Localization of Transcription Factors Downstream of DAF-2/IIS

When DAF-2/IIS is activated, both DAF-16/FoxO and SKN-1/Nrf are phosphorylated and excluded from the nucleus; however, upon down-regulation of DAF-2/IIS, both transcription factors are shuttled into the nucleus (Ogg et al. 1997; Tullet et al. 2008). To observe any alterations to this translocation, we crossed DAF-16::GFP and SKN-1::GFP transgenes into a *dbl-1* mutant background and treated animals with *daf-2* RNAi. Localization of the two transcription factors was observed in the intestinal nuclei due to their size and prominence.

We categorized nuclear localization of the GFP fusion proteins in these backgrounds as none (cytoplasmic only), low (diffuse in both cytoplasm and nucleus), medium (faint nuclear localization) or high (strong nuclear localization). In a wild-type background, animals treated with the empty vector, L4440, showed DAF-16::GFP fluorescence diffusely or weakly nuclear, with a small percentage showing only cytoplasmic localization (Figure 3). As expected, treatment with *daf-2* RNAi caused nuclear accumulation of DAF-16::GFP in all animals (p = 0.0017). In *dbl-1* mutant animals on control RNAi plates, there was no significant difference in the level of nuclear localization compared with the wild-type controls (Figure 3). Upon reduction of *dbl-1* and *daf-2* together, however, intensity of nuclear DAF-16::GFP became higher than in *daf-2(RNAi)* animals alone (p < 0.0001) (Figure 3). We conclude that loss of *dbl-1* increases the responsiveness of the DAF-16::GFP reporter to DAF-2 activity.

**Figure 3.**
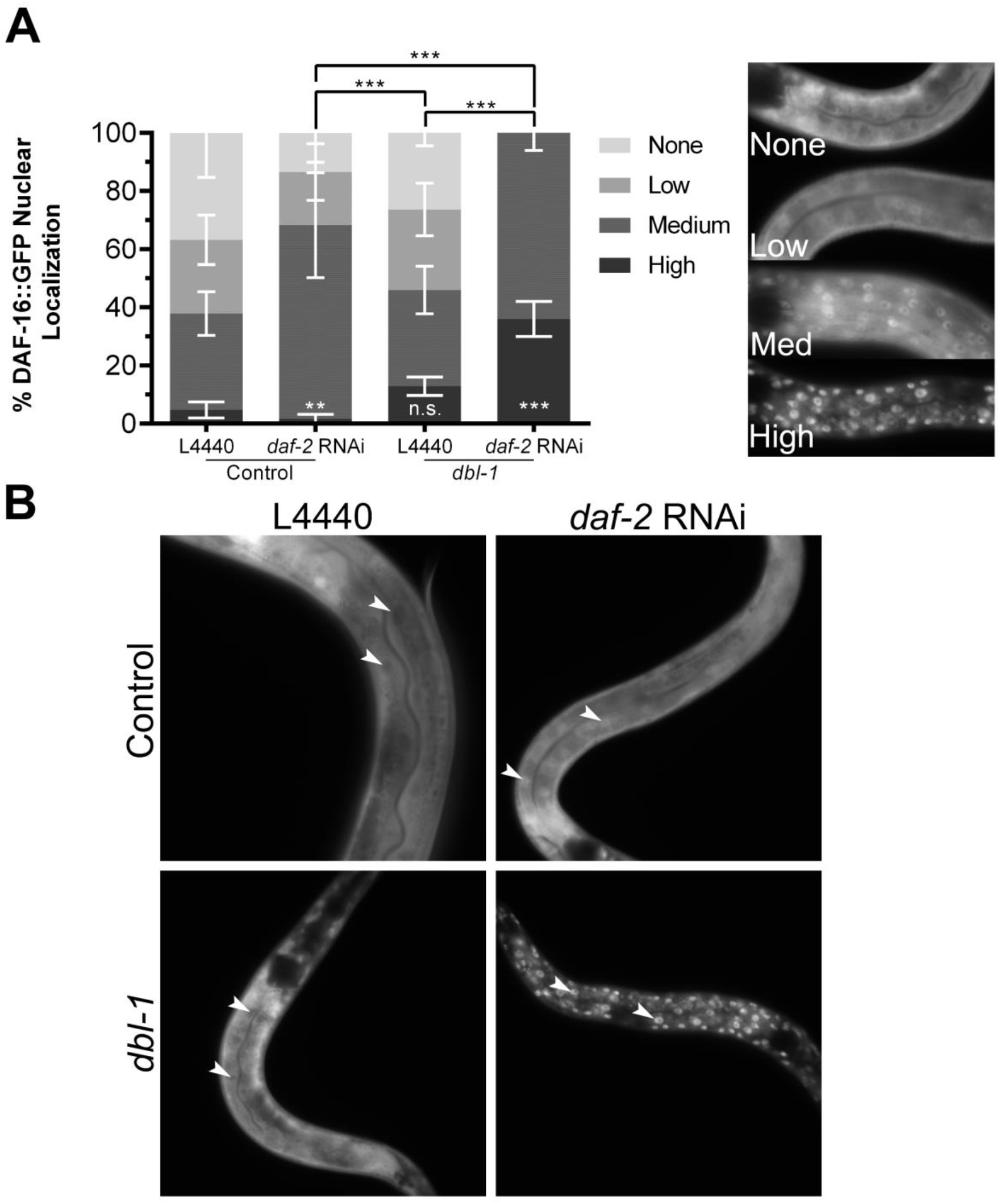
DAF-16 localization is altered in a *dbl-1* mutant background. Control animals treated with empty vector exhibit some nuclear localization of DAF-16::GFP, upon treatment with *daf-2* RNAi, DAF-16::GFP nuclear localization increases. In *dbl-1* mutants, DAF-16::GFP nuclear localization is not significantly changed compared to the control under empty vector treatment. Upon treatment with *daf-2* RNAi, a higher level of nuclear localization is observed as compared to control animals treated with *daf-2* RNAi. Asterisks across the bottom represent significance compared to Control, n.s. not significant, ** p<0.01, *** p<0.001, error bars represent SEM. B) Images of animals expressing DAF-16::GFP taken at 400X. White arrows indicate examples of intestinal nuclei. Camera settings were identical between all strains.

In a wild-type background, animals treated with the empty vector, L4440, exhibited mainly diffuse expression of SKN-1::GFP, with very few animals containing nuclear localization. Upon treatment with *daf-2* RNAi, the majority of animals showed a significant increase in nuclear SKN-1::GFP, as expected (p<0.0001). When treated with L4440, *dbl-1* mutants exhibited an increased level of nuclear localization compared to the wild-type control (p=0.0073), but less than that of wild type treated with *daf-2* RNAi (p<0.0001). Interestingly, upon treatment with *daf-2* RNAi, localization of SKN-1::GFP in *dbl-1* mutants was not significantly different from those treated with the empty vector (p=0.8694) (Figure 4A,B). This outcome contrasts with the results with DAF-16::GFP. Instead, the responsiveness of the SKN-1::GFP reporter to the reduction of DAF-2/IIS was reduced in the *dbl-1* mutant background.

**Figure 4.**
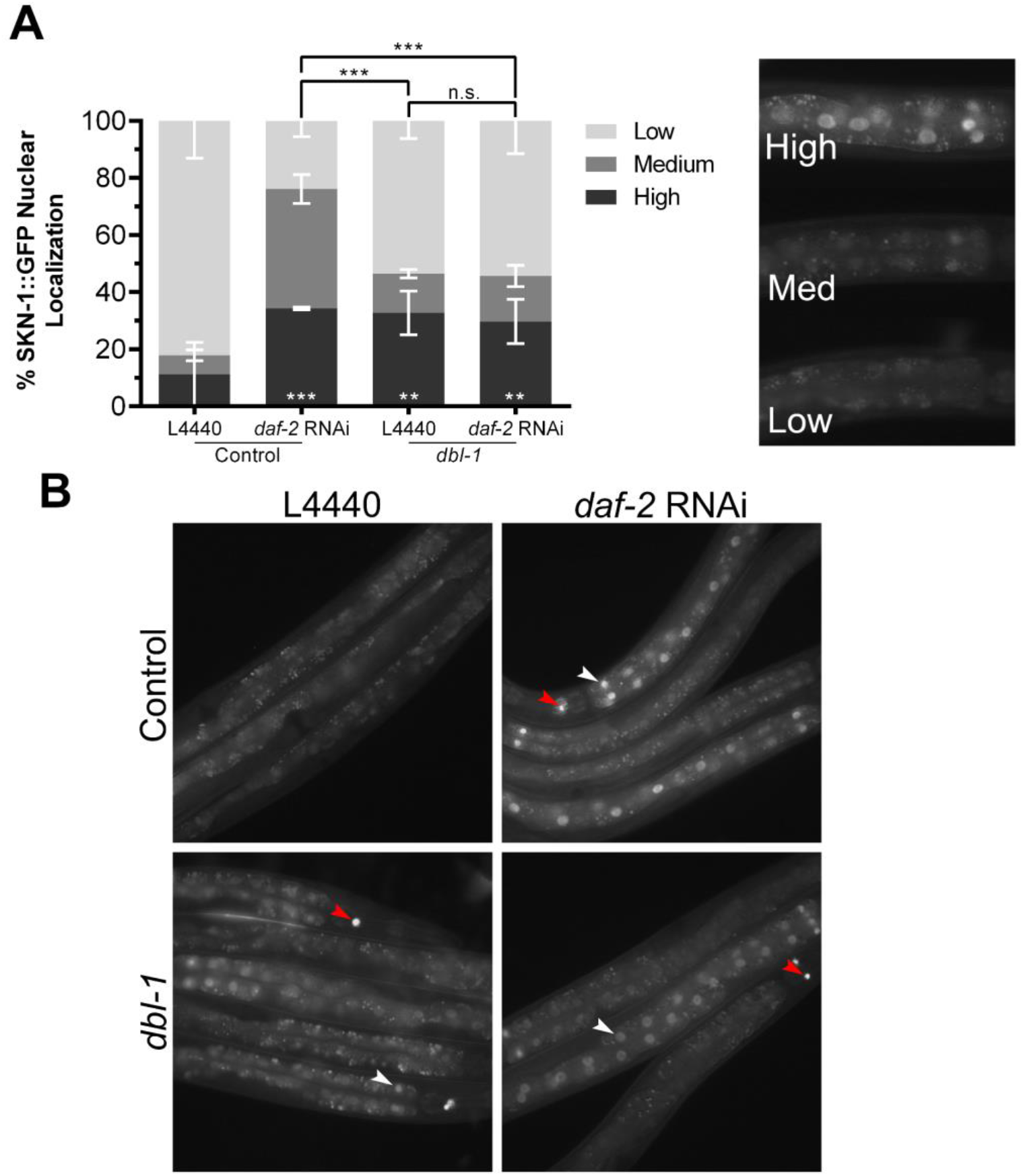
SKN-1::GFP localization is altered in *dbl-1* mutants regardless of DAF-2/IIS. A) Control animals treated with empty vector exhibit very little nuclear localization of SKN-1::GFP, upon treatment with *daf-2* RNAi, SKN-1::GFP nuclear localization drastically increases. In *dbl-1* mutants, SKN-1::GFP nuclear localization is slightly increased compared to the control under empty vector treatment, upon treatment with *daf-2* RNAi, no change in nuclear localization is observed. Nuclear localization was graded as high, medium, or low, with an example of each ranking shown. Asterisks across the bottom represent significance compared to Control, n.s. not significant, ** p<0.01, *** p<0.001, error bars represent SEM. B) Images of animals expressing SKN-1::GFP taken at 200X. White arrows indicate examples of nuclei, red arrows indicate SKN-1::GFP expression in ASI neurons. Camera settings were identical between all strains.

### Effects of IIS Transcription Factors on *dbl-1* Low-Fat Phenotype

We sought to determine whether one or both of these transcription factors mediate regulation of fat storage downstream of DBL-1/BMP and DAF-2/InsR. The strains expressing tagged functional DAF-16/FoxO or SKN-1/Nrf contain multicopy arrays that may cause an increase in the expression levels of these transcription factors. We measured fat accumulation using Oil Red O staining in L4 stage animals. In the wild-type background, neither *daf-16::gfp* nor *skn-1::gfp* have increased fat stores (Figure 5). Next, we examined *dbl-1* mutant animals. As expected, *dbl-1* mutants have low fat stores. *daf-16::gfp* expression, but not *skn-1::gfp* expression, is capable of suppressing the low-fat phenotype of *dbl-1* mutants (observed in two out of three trials). This result suggests that SKN-1 does not mediate the effect of DBL-1 on fat accumulation.

**Figure 5.**
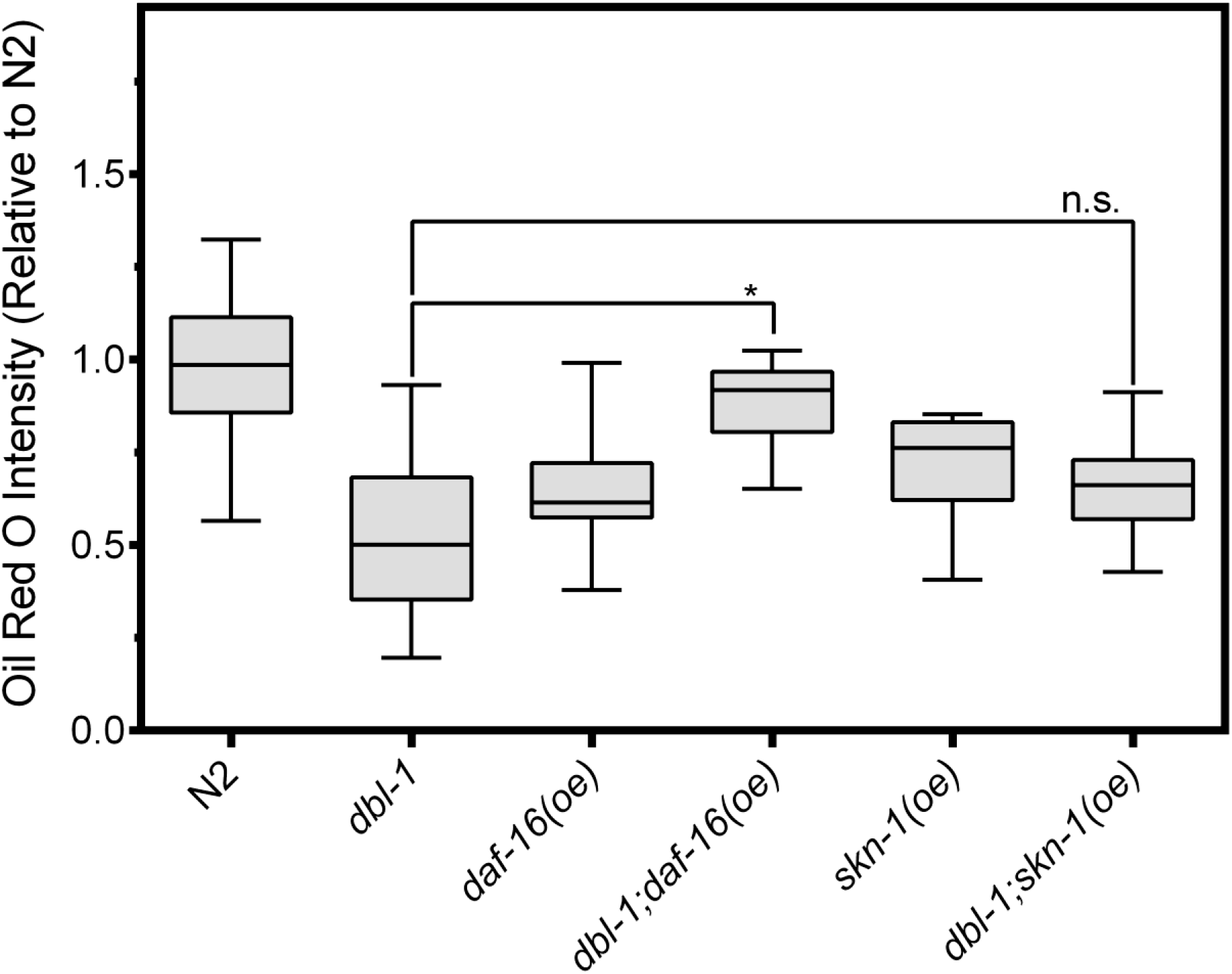
Effects of DAF-16/FoxO and SKN-1/Nrf overexpression on lipid accumulation. Animals were stained at L4 stage with the neutral lipid dye Oil Red O. Images taken at 400X, camera settings were kept constant between all strains. Quantification was done for equivalent regions of the intestine just posterior to the pharynx. n.s. not significant, * p value < 0.05, *** p value < 0.001, boxes denote the 2^nd^ and 3^rd^ quartiles, whiskers denote min and max values.

## Discussion

We previously showed that DBL-1/BMP signaling and DAF-2/IIS interact to regulate lipid stores in *C. elegans* (Clark et al. 2018). In this study, we identify additional interactions by presenting evidence that the DBL-1/BMP signaling pathway exhibits an antagonistic effect on dauer formation and autophagy induction in a DAF-2/InsR sensitized mutant background. We show that loss of BMP signaling activity causes a reduction in the rate of both dauer formation and autophagy, in *daf-2* mutants at 20°C. Conversely, increased BMP signaling causes an increase in the rates of these phenotypes. These interactions are only revealed in a DAF-2-compromised background, distinguishing them from the epistatic relationship of BMP and IIS in lipid regulation and the independent functions of BMP and IIS in body size regulation and reproductive aging (Luo et al. 2010; Clark et al. 2018). Our findings implicate coordination between BMP and IIS pathways in mutliple aspects of development and homeostasis, an interpretation that is supported by additional evidence in both *C. elegans* and other systems. For example, DBL-1/BMP interacts with IIS in *C. elegans* L1 arrest. In *C. elegans*, embryos that hatch without food enter an L1 arrest in which cell divisions are blocked. The arrested somatic development of different cell lineages has been shown to be dependent on DBL-1/BMP interaction with IIS. In mesodermal lineages, the cell division arrest is defective in *daf-16* mutants, and DBL-1/BMP activity is required for the L1 arrest defective phenotype of *daf-16* (Kaplan et al. 2015). In neuronal lineages, DBL-1/BMP acts upstream to promote IIS activity in a DAF-16-independent role, through reduction in the expression of *ins-3* and *ins-4* in neurons (Zheng et al. 2018b). In addition, DBL-1/BMP signaling interacts with DAF-16/FoxO function to induce germline tumor formation (Qi et al. 2017). In vertebrates, tissue culture studies have shown that BMP2 and BMP6 regulate insulin sensitivity in adipose cells (Schreiber et al. 2017). In mice, BMP4 regulates insulin sensitivity of adipose tissue (Qian et al 2013.). Furthermore, BMP7 regulates expression of insulin signaling components in C3H10T1/2 cells, leading to fat browning (Zhang et al. 2010). Numerous additional instances of interactions between TGFβ/activin signaling and IIS also exist, emphasizing the widespread deployment of this signaling axis.

This work has uncovered specific components of the IIS pathway that mediate the effects of DBL-1/BMP on lipid accumulation. At least two ILPs, INS-4 and INS-7, are transcriptionally repressed by DBL-1/BMP signaling (Liang et al. 2007). Using an *ins-4p::gfp* reporter, we now show that *ins-4* expression is highly and specifically increased in the hypodermis in a *sma-3* mutant background, indicating that Smad signaling regulates *ins-4* expression in the hypodermis. Additionally, we show that *ins-4* mutants have increased levels of lipids, demonstrating that, in spite of any potential redundancy between ILPs, INS-4 is critically required for regulation of lipid storage. Consistent with our analysis, overexpression of *ins-4* is sufficient to lower fat levels (Zheng et al. 2018a). These data provide evidence that DBL-1 activates Smad signaling in the hypodermis, down-regulating the expression of INS-4, to reduce DAF-2/IIS, in turn, maintaining proper lipid stores in the intestine (Figure 6). The other ILP gene we identified as a transcriptional target of DBL-1, *ins-7*, has no effect on fat accumulation when overexpressed (Zheng et al. 2018a). Since *ins-7* is a known transcriptional target of IIS in the intestine (Murphy et al. 2007), its upregulation in *dbl-1* mutants may be a secondary consequence of increased IIS activity. We also interrogated the effects of downstream IIS transcription factors DAF-16/FoxO and SKN-1/Nrf on DBL-1-mediated fat regulation. Since these transcription factors act in opposition to the other IIS components, we did not analyze loss-of-function mutants but rather overexpression from multi-copy transgenes. Our results are consistent with DAF-16/FoxO, but not SKN-1/Nrf, mediating the effect of DBL-1 on fat accumulation (Figure 6).

**Figure 6.**
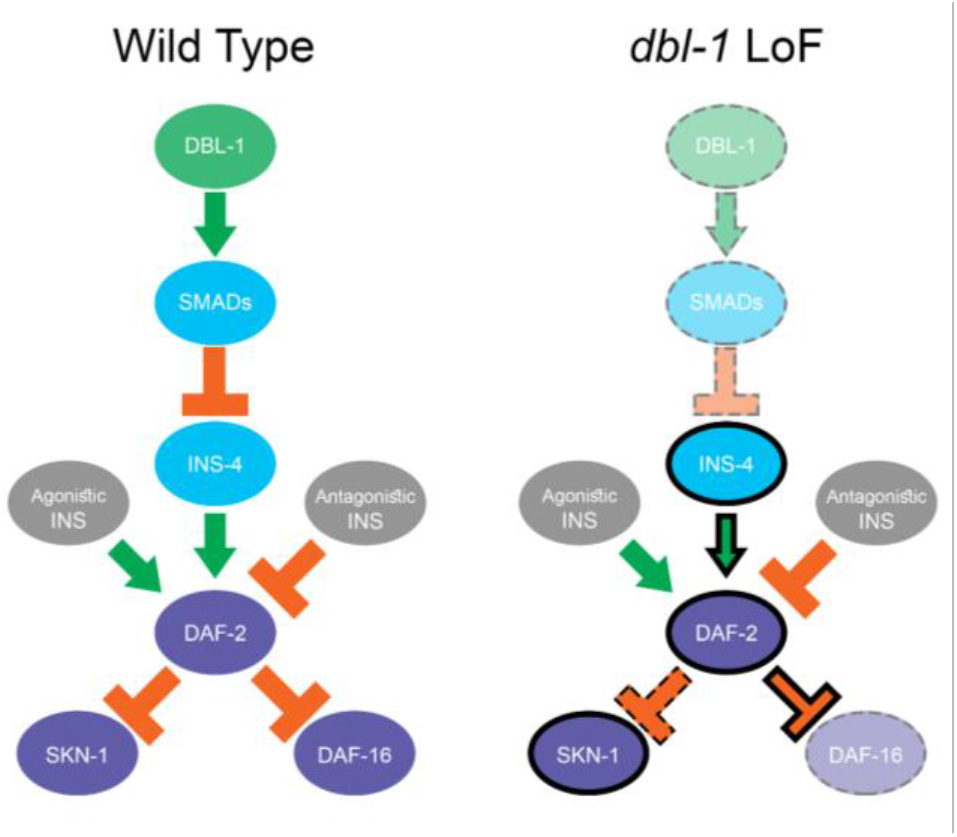
Model for interaction between DBL-1/BMP signaling and DAF-2/IIS pathways. In wild-type animals, DAF-2/InsR is regulated by multiple agonistic and antagonistic ILPs. DAF-2/InsR activation leads to phosphorylation and cytoplasmic retention of SKN-1/Nrf and DAF-16/FoxO. In *dbl-1* loss-of-function (LoF) mutants, reduced Smad activity leads to increased INS-4 expression. Also in *dbl-1* mutants, SKN-1/Nrf nuclear localization has reduced sensitivity to DAF-2/InsR activity, while DAF-16/FoxO nuclear localization has increased sensitivity to DAF-2/InsR function.

IIS transcription factors DAF-16/FoxO and SKN-1/Nrf are regulated, in part, at the level of subcellular localization. DAF-2/IIS activity promotes cytoplasmic retention of DAF-16/FoxO and SKN-1/Nrf, inhibiting their functions as transcription factors. In this study, we find that DBL-1/BMP activity alters the sensitivity of DAF-16/FoxO and SKN-1/Nrf to DAF-2/IIS in opposite ways. In *dbl-1* mutants, SKN-1::GFP localization is not significantly altered by *daf-2* RNAi (Figure 2), indicating that DBL-1/BMP is required for full responsiveness of SKN-1/Nrf to DAF-2/IIS. For DAF-16::GFP, we see the opposite effect on sensitvity to DAF-2/IIS. In *dbl-1* mutants, DAF-16::GFP is less nuclear than control and upon treatment with *daf-2(RNAi)* it becomes intensely nuclear (Figure 3). If DBL-1/BMP signaling caused a simple reduction in DAF-2/IIS activity, we would have expected both transcription factors to respond similarly, with increased cytoplasmic localization in a *dbl-1* mutant and possibly equivalent nuclear localization upon *daf-2* RNAi in both wild-type and *dbl-1* backgrounds. Two possible mechanisms for the observed differences between DAF-16 and SKN-1, which are not mutually exclusive, are that INS-4 activation of DAF-2/InsR differentially affects the downstream transcription factors, or that DBL-1/BMP signaling has an additional direct influence on localization independent of IIS.

While specific domains of expression can provide fine-tuned control of ligand-receptor combinations in an organism, individual ligands can also produce alternate responses through a single receptor. In mammals, the Fibroblast Growth Factor (FGF) signaling family consists of 18 ligands (FGFs) and four tyrosine kinase receptors (FGFRs) (Ornitz and Itoh 2015). In the FGF signaling family, there exist examples of two ligands with overlapping expression patterns that elicite different responses from the same receptor. Both FGF7 and FGF10 exhibit roles in the branching morphogeneis of the lacrimal and submandibular glands in mice using *ex vivo* organ cultures. These two ligands both bind to FGFR2b, however, FGF7 induces a proliferative response while FGF10 induces a migratory response, switching morphogenesis between branching and elongation respectively (Steinberg et al. 2005; Makarenkova et al. 2009). The mechanism thought to underly this switch is through the differential phosphorylation of a specific tyrosine residue. Binding of FGF10 to FGFR2b results in the phosphorylation of residue Y734, which recruits a complex involved in endocytic recycling, while binding of FGF7 does not (Francavilla et al. 2013). The exact mechanism by which the two ligands result in differential phosphorylation is unknown, however structural evidence suggests that binding of the ligands may induce two different confirmations of the intracellular domains (Bae and Schlessinger 2010). While this mechanism has not been studied in the context of DAF-2/IIS in *C. elegans*, it does not preclude the possibility that this mechanism may be shared between the two tyrosine kinase receptor families. While the 40 insulin-like peptides present in *C. elegans* can be classified as either agonists or antagonists of DAF-2, differential effects within a classification could be attributed to a similar mechanism.

Altered subcellular localization of DAF-16/FoxO provides a mechanism for the demonstrated inhibitory interaction between BMP and IIS in lipid metabolism. In addition to the interactions already discussed, Qi et al. identified a cooperative interaction between DBL-1/BMP signaling and DAF-16/FoxO that does not occur at the level of subcellular localization (Qi et al. 2017). They find that DAF-16/FoxO functions nonautonomously in the hypodermis to induce germline tumor formation, and that this function depends on DBL-1/BMP signaling. This phenotype is independent of nuclear localization of DAF-16/FoxO; rather the Smads SMA-2 and SMA-3 were shown to bind DAF-16 directly and presumably promote its transcriptional activity (Qi et al. 2017). Taken together, these analyses exemplify the interconnected, polymodal nature of BMP and IIS pathway interactions in the regulation of organismal function. The homeostatic regulation of cellular processes, such as metabolism, are vital to the well-being of an organism and must be flexible enough to coordinate a variety of inputs and outputs. In mammals, the complexity of tissues involved in such crosstalk creates a challenge for precise analysis of these interactions. Using *C. elegans*, we have developed a model to understand the relationship of two highly conserved pathways, BMP and IIS, in the intact organism.

## Acknowledgments

We thank Michael Meade and Shoshana Reich for the construction of CS663 and CS633, respectively. We thank Nick Palmisano for input on autophagy assays. This work was supported in part by National Institutes of Health R15GM112147 to CSD, and by 2R15GM102846-02 to AM. Some strains were provided by the CGC, which is funded by NIH Office of Research Infrastructure Programs (P40OD010440). This work was carried out in partial fulfillment of the requirements for the Ph.D. degree from the Graduate Center of City University of New York (JFC and EJC).

---

**S1 Fig.**
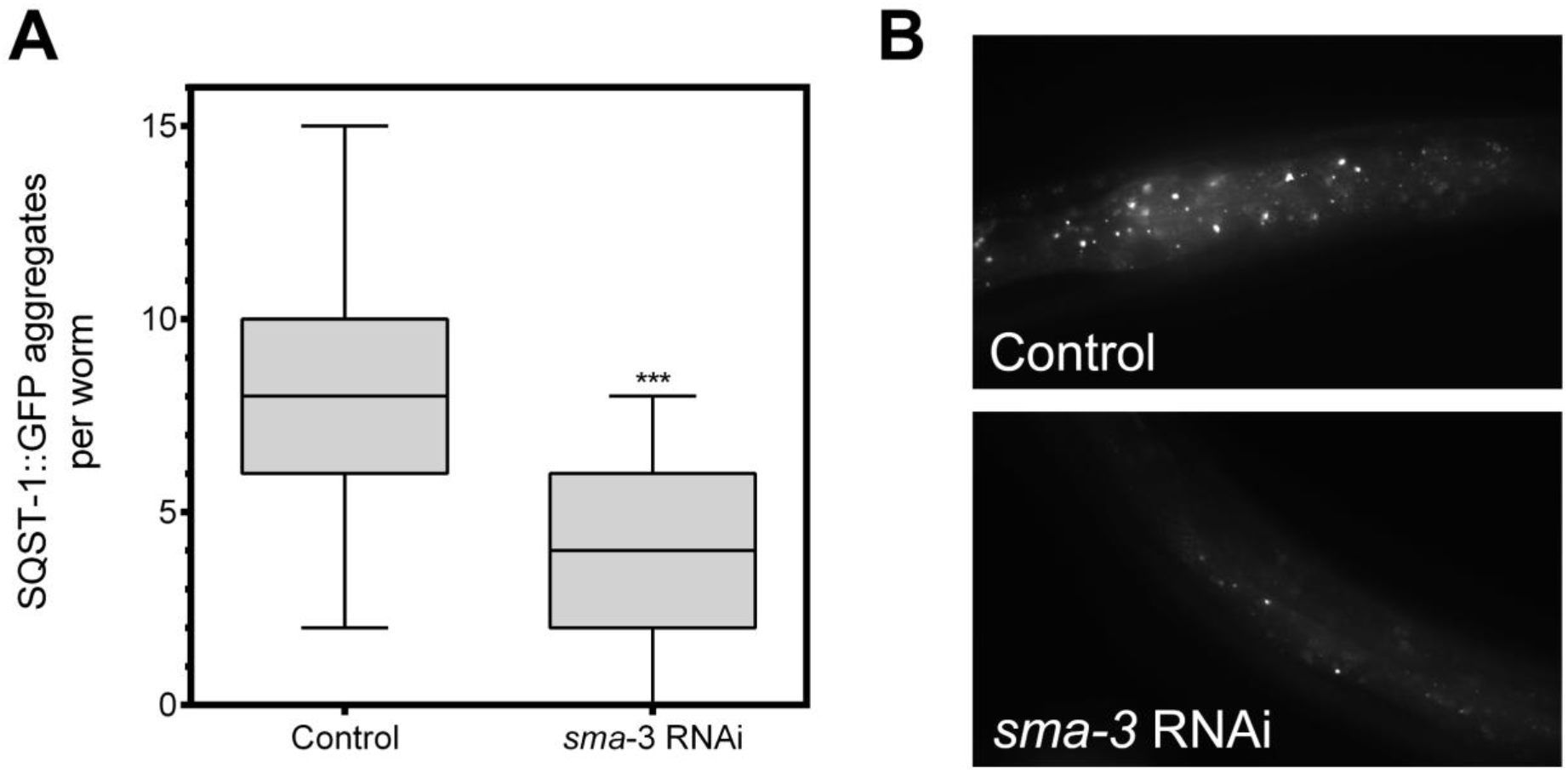
Inactivation of *sma-3/Smad* suppresses the accumulation of SQST-1::GFP aggregates in *rpl-43* mutants. A) *rpl-43* mutants show accumulation of SQST-1::GFP puncta due to impaired autophagic flux (Guo et al. 2014). Genetic backgrounds that increase autophagy allow the clearance of those aggregates. We analyzed this strain at the L4 stage following RNAi depletion of *sma-3/Smad* compared with empty vector control (L4440), and found a significant clearance of SQST-1::GFP puncta (p < 0.001) as expected (Guo et al. 2014). Two trials of n = 15 animals yielded similar results, and the combined data are shown. B) Images of *sqst-1::gfp* animals treated with either an empty vector control (L4440) or *sma-3* RNAi.

